# Positional identification of a candidate gene for *MALE STERILITY 2* (*MS2*) by linkage mapping and transcriptomic data in *Cryptomeria japonica* D. Don

**DOI:** 10.1101/2025.09.11.674818

**Authors:** Saneyoshi Ueno, Yoichi Hasegawa, Tokuko Ujino-Ihara, Momi Tsuruta, Hiroyuki Kakui, Junji Iwai, Satoko Hirayama, Katsushi Yamaguchi, Shuji Shigenobu, Takeshi Fujino, Yutaka Suzuki, Masahiro Kasahara, Yoshinari Moriguchi

**Author notes:** Corresponding authors: Saneyoshi Ueno, Department of Forest Molecular Genetics and Biotechnology, Forestry and Forest Products Research Institute, Forest Research and Management Organization, Matsunosato, Tsukuba, Ibaraki, 305-8687, Japan; Yoshinari Moriguchi, Department of Agriculture, Faculty of Agriculture, Niigata University, 8050 Ikarashi 2-no-cho, Nishi-ku, Niigata, 950-2181, Japan. Junji Iwai, Forestry Conservation Division, Department of Agriculture, Forestry and Fisheries, Niigata Prefectural Government, Niigata, Japan; Satoko Hirayama, Forestry Administration Division, Department of Agriculture, Forestry and Fisheries, Niigata Prefectural Government, Niigata, Japan.

## Abstract

**Background:** Japanese cedar (*Cryptomeria japonica* D. Don) is a major plantation species in Japan, but its abundant pollen production is a primary cause of seasonal allergic rhinitis (pollinosis). To mitigate this public health issue, the use of male-sterile cultivars has been promoted. Five types of recessive male-sterile mutations (*ms1*–*ms5*) have been identified, and the causal genes and mutations for *MS1* and *MS4* have been elucidated. However, the gene responsible for MS2-type male sterility remains unknown.

**Results:** We aimed to identify the candidate gene responsible for MS2-type male sterility using a map-based cloning strategy. High-resolution linkage mapping localized the *MS2* locus to a 1.56 cM interval on linkage group 5, corresponding to an 8.64 Mb region of the reference genome. Ninety-one genes in this region were subjected to functional annotation, gene expression analysis, and mutation screening. Among these, a single gene, *SUGI_0493010*, encoding a GDSL-type esterase/lipase protein (GELP), fulfilled all three criteria: it showed homology to pollen development genes in *Arabidopsis thaliana*, was specifically expressed in male strobili, and carried a deleterious amino acid substitution (S40F) within the predicted catalytic domain in *ms2* mutant. The same mutation was also detected in a heterozygous individual (*Ms2/ms2*) from a separate breeding population, whose genotype was confirmed through progeny testing. Structural annotation revealed that the affected serine residue lies within the conserved GDSL motif, suggesting a functional disruption of enzymatic activity.

**Conclusions:** Our results strongly suggest that SUGI_0493010 (*GELP*) is the causal gene for MS2-type male sterility in *C. japonica*. This finding enhances our understanding of male sterility mechanisms in conifers and provides a valuable genetic resource for breeding pollen-free trees. The study also demonstrates the effectiveness of combining genetic mapping with transcriptomic and mutational data in forest tree genomics.

## Introduction

Japanese cedar (*Cryptomeria japonica* D. Don), or sugi, is one of the most widely planted tree species in Japan due to its economic importance in the forest industry, occupying about 40 % of artificial forests in Japan [1]. The timber has been used not only in house construction, but also in traditional crafts and daily-use wooden products, including furniture, chopsticks, sake barrels, and shrine architecture. However, large-scale sugi plantations have contributed to a major public health issue: pollen allergy or sugi pollinosis. This seasonal allergy has become increasingly prevalent, with nationwide surveys indicating that nearly 40% of the population was affected in 2019 [2]. The primary source of the allergen is the massive production of pollen from male strobili during the spring season, varying widely among studies, ranging from approximately 100,000 to 656,000 grains per single male strobilus depending on the clone and environmental conditions [3–6]. In order to reduce pollen emissions at the source, increasing attention has been directed toward the utilization of male-sterile cultivars in afforestation.

Genetic characterization of male-sterile mutants in *C. japonica* has provided a solid foundation for this approach. Recessive mutations causing male sterility have been classified into five types (MS1–MS5) based on the specific developmental stage at which pollen formation is disrupted. These classifications were established through detailed cytological observations [7–10] and linkage analyses using controlled crosses [11–13]. Among these, the MS1-type has been the most extensively studied [8, 14, 15] and has already been utilized in public afforestation initiatives [16]. Pollen-free sugi trees homozygous for the *ms1* allele (*ms1/ms1*) have been propagated and planted to reduce airborne pollen emission from sugi forests. Recent molecular studies [17, 18] have further advanced our understanding of the genetic basis of male sterility in *C. japonica*, particularly for *MS1* and *MS4*. *MS1*, which is associated with complete pollen abortion at the tetrad stage, has been linked to a gene involved in lipid transportation. Similarly, *ms4*, which disrupts pollen formation during the microspore stage, has been associated with a gene orthologous to *Arabidopsis thaliana TKPR1*, a key enzyme in sporopollenin biosynthesis [19]. These findings have provided valuable insights into the molecular mechanisms underlying pollen development and laid the foundation for marker-assisted selection of male-sterile trees in breeding programs [20, 21].

In contrast, the molecular basis of MS2-type male sterility remains unresolved. *MS2* is characterized by developmental abnormality from the tetrad stage, as previously reported in [10] (Figure 1) and was first identified in a naturally occurring mutant known as ‘Shindai-1,’ selected from an artificial forest in Niigata Prefecture in 1998 [9]. Previous studies have localized the *MS2* locus to linkage group 5 (LG5) [11, 22], but the causal gene has not yet been identified. Understanding the genetic mechanism underlying *MS2* is essential not only for expanding the tools for marker-assisted breeding but also for deepening our knowledge of reproductive development in conifers.

**Figure 1.**
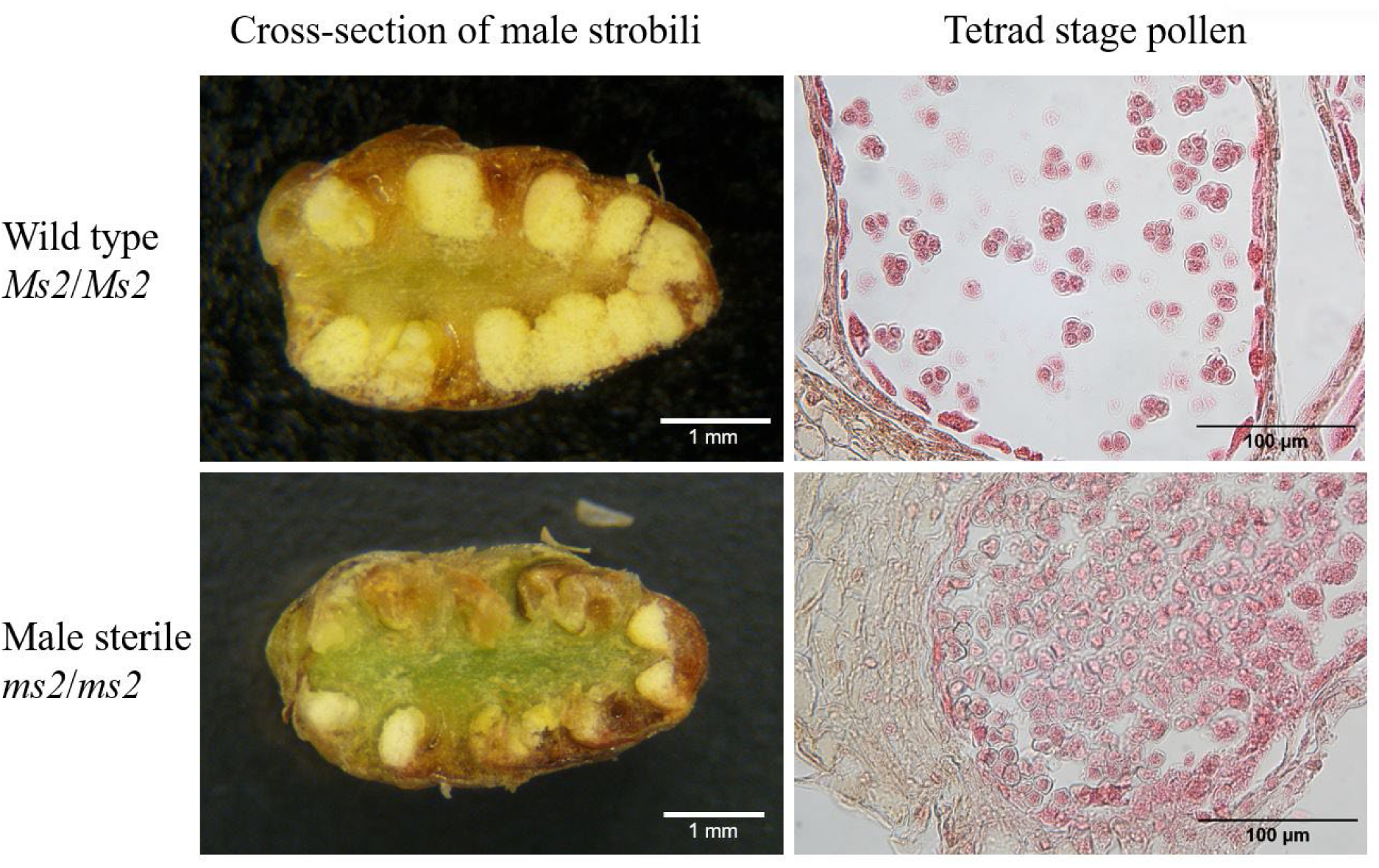
Comparison of male strobili and tetrad stage pollen between fertile and MS2-type male-sterile individuals of *Cryptomeria japonica*. Representative images showing the morphological and cytological differences between wild-type (*Ms2*/*Ms2*) and MS2-type male-sterile (*ms2*/*ms2*) individuals. **Left**: Cross-sections of male strobili. **Right**: Light microscopic image of pollen at the tetrad stage. In fertile individuals, pollen development proceeds normally, while in sterile individuals (‘Shindai-1’), tetrads exhibit developmental abnormality with clumping and dense aggregation. Scale bars for the cross-sections are estimated based on the approximate size of the male strobilus (∼4 mm) and are intended for reference only.

In this study, we aimed to identify the candidate gene responsible for MS2-type male sterility using a map-based cloning approach. By integrating high-resolution linkage mapping, RNA-Seq-based expression profiling, and mutation screening with functional prediction, we successfully narrowed down the candidate region and identified a GDSL-type esterase/lipase gene (*GELP*) as the most likely causal gene. Our findings provide important insights into the molecular basis of male sterility in *C. japonica* and offer a genetic resource for the practical development of pollen-free sugi trees.

## Materials and Methods

### Plant materials

The MS2-type male-sterile individual ‘Shindai-1’ was originally selected from an artificial forest in Niigata Prefecture, Japan [9]. This individual was used as the seed parent to produce a mapping family (S1-2), which was generated by crossing ‘Shindai-1’ (*ms2*/*ms2*) and ‘S1NK4’ (*Ms2*/*ms2*) as female and male parent, respectively [22]. The ‘S1NK4’ is an offspring (F1) resulting from a cross between ‘Shindai-1’ and ‘Nakakubiki-4.’ Among the offspring of this family, 129 individuals were used for refinement of the linkage map in our previous study [22]. For the fine mapping of the *MS2* locus, we selected 95 of these 129 individuals (31 male-sterile and 64 male-fertile). Because the *MS2* male-sterile phenotype can be difficult to distinguish in some individuals, only those with clearly identifiable phenotypes were included in the analysis. Phenotypes were determined based on field observations of male strobili and microscopic examination of dissected male strobili. In addition, another individual from Niigata Prefecture, ‘Gosenshi-1,’ which had been previously identified as *Ms2/ms2* through controlled crossing experiments [23], was included to evaluate the consistency of the candidate mutation identified in this study.

### RNA-Seq data collection and analysis

Male strobili were collected from fertile and sterile individuals of the S1-2 family growing at Chiyoda nursery, FFPRI (36.184269° N, 140.217564° E) at two time points (October 17th and November 6th, 2019) for RNA-Seq analysis (Additional file 1). The male strobilus samples were immediately frozen in liquid N_2_ at the field, and stored at −80°C until RNA extraction. The total RNA was extracted from the male strobili using the modified cetyltrimethylammonium bromide (CTAB) method [24]. RNA-Seq was performed following Macrogen’s regular workflow (Macrogen Japan Corp., Kyoto, Japan). Briefly, high-throughput sequencing libraries were constructed using the TruSeq stranded mRNA Library Kit (Illumina, San Diego, CA, USA) following the quality control of RNA. Each library was sequenced by the Illumina platform (NovaSeq6000) at 100 bp paired-end reads (on average 7.3 Gbp per sample). Raw reads were processed using fastp [25] to remove adapter sequences and low-quality regions. Cleaned reads were aligned using bwa mem (ver. 0.7.17-r1188) with default parameters. Depending on the timing of data generation, either the *Cryptomeria japonica* (SUGI_1) permissive gene set [26] or the earlier transcript reference CJ3006NRE [27] was selected as the mapping reference. BAM files aligned to the SUGI_1 permissive gene set were subsequently utilized as a common resource for downstream analyses—including SNP identification, marker development, gene expression analysis (including DEG analysis), and detection of deleterious mutations—all of which focused on the SUGI_1 standard gene set [26] consisting of 55,246 annotated genes.

### Marker development using RNA-Seq data

Based on our previous linkage map [22], the *MS2* locus was initially localized to three genomic contigs—ctg668, ctg930, and ctg820—in the *C. japonica* draft genome ver. 0.1 (Fujino et al., unpublished) (Additional file 2). At that time, a chromosome-scale genome assembly and gene annotations were not yet available. Therefore, we identified genes on these contigs by performing BLASTN searches using CJ3006NRE reference transcript sequences from our previous study [27] as queries. To improve the accuracy of gene models and prioritize polymorphisms, spliced alignments were performed using Splign [28].

Polymorphisms between fertile and sterile individuals were identified using RNA-Seq data. BAM files were grouped by phenotype (fertile or sterile), and SNPs were called using VarScan (v2.4.3) [29, 30] with default parameters, generating VCF files for downstream analysis. Within each target gene, consensus sequences incorporating heterozygous sites were created using ‘bcftools consensus’ (v1.9) [31] with IUPAC ambiguity codes. These sequences were then post-processed to fit the input format of Fluidigm SNPType Assay software D3 (Standard Biotools).

### SNP genotyping and fine mapping

Candidate SNPs were evaluated using D3 to determine whether functional primer pairs could be designed. Only SNPs that passed the D3 design criteria were selected for genotyping. Genotyping was carried out using the Fluidigm EP1 system with a 48.48 Dynamic Array according to the manufacturer’s instructions. Genotypes were called with Fluidigm SNP Genotyping Analysis software (v4.5.1). The resulting genotype data were used to construct a refined linkage map. Linkage analysis was conducted using the maximum likelihood mapping algorithm in JoinMap ver. 5 (Kyazma, Wageningen, The Netherlands), assuming a backcross (BC)-type population (nn × np), as previously described [22]. Once a chromosome-scale genome assembly (SUGI_1) became available [26], the newly developed markers were mapped onto chromosome 5 (chr5).

### Candidate gene identification by functional annotation

To identify candidate genes within the *MS2* region described above, nucleotide sequences corresponding to the standard gene set [26] in this region (Additional file 3) were subjected to BLASTX searches (E-value cutoff: 1e-5) against the TAIR10 protein database of *Arabidopsis thaliana* [32]. Genes in *C. japonica* showing homology to 372 *Arabidopsis* gene loci annotated with GO:0009555 (pollen development) were selected as functional (homology-based) candidates.

### Gene expression analysis for candidate gene prioritization

Gene expression analysis was performed to identify candidate genes expressed in fertile individuals within the 8.64 Mb interval on chromosome 5. Gene expression levels were quantified using transcripts per million (TPM) values calculated from the RNA-Seq data using featureCount [33], based on BAM files aligned to the SUGI_1 permissive gene set. TPM values were calculated across all annotated transcripts, and only genes corresponding to the SUGI_1 standard gene set (55,246 genes) were retained for downstream analysis. Genes with TPM values > 1 in at least one fertile individual were considered expressed and retained as MS2 candidate genes. Differential gene expression analysis was conducted using DESeq2 [34] with raw count data from the SUGI_1 permissive gene set. For interpretation, we focused on the MS2-neighboring genes. Genes with an adjusted *p*-value < 0.01 were considered significantly differentially expressed.

### Identification of deleterious mutations and validation in additional material

To investigate whether deleterious mutations were responsible for MS2-type male sterility, polymorphisms detected within candidate genes were evaluated using PROVEAN (v1.1.3) [35]. Variant information from the previously generated VCF file was used to create sterile (ALT) haplotype sequences by using ‘bcftools consensus’ (ver. 1.9) [31], translated into protein sequences by transeq in EMBOSS suite [36], and the resulting amino acid substitutions were analyzed for their functional effects using PROVEAN (v1.1.3). Variants with a PROVEAN score below –2.5 were classified as deleterious. Candidate genes with deleterious mutations were prioritized for further investigation.

To further validate the presence of the identified mutation in an independent genetic background, we also analyzed an additional heterozygous individual, ‘Gosenshi-1’ (*Ms2*/*ms2*), from which male strobili were collected at Niigata Prefectural Forest Research Institute on 22 October 2020 for RNA extraction. RNA extraction and mutation detection for ‘Gosenshi-1’ were performed using the same methods as described above.

### Structural, functional, and expression profiling of candidate gene

The gene structure of a candidate gene (SUGI_0493010) was determined using the SUGI_1 genome assembly and gene models from the standard gene set [26]. To refine transcriptional boundaries, full length cDNA sequence (DDBJ accession number: FX344585) and Iso-Seq reads for male strobili from our previous studies [27, 37] were mapped using minimap2 [38] with options (-ax splice -uf -k14 --secondary=no) and visualized in CLC Genomics Workbench ver. 20 (Qiagen). Transcription start site (TSS) was inferred based on sharp increases in read coverage, and on the prediction by TSSPlant [39] with input of a 3.4-kb upstream sequence from the start codon. The 5′UTR was defined as the region between the TSS and the annotated start codon, while the 3′UTR was defined as the region between the stop codon and the major polyadenylation site inferred from Iso-Seq read ends. The predicted amino acid sequence (364 residues) was annotated using InterPro webtool [40].

In order to get insights into expressional variation among different tissues for the MS2 candidate gene (SUGI_0493010), CPM (count per million) data were downloaded from the SugiExDB database [41], which contains RNA-Seq expression profiles for various *C. japonica* tissues. Expression levels were visually compared across tissues to explore tissue-preferential expression tendencies.

## Results

### Fine mapping of the *MS2* locus

To refine the localization of the *MS2* gene, we constructed a high-resolution linkage map using the mapping population. The locus was positioned between two flanking markers, CJt005282-890 and CJt113083-201, spanning an 8.64 Mb region on chromosome 5 (Figure 2 and Additional file 2). RNA-Seq-derived markers served as anchor markers, bridging the genetic linkage map and the physical genome map, and enabled accurate localization of genes within the MS2 locus and supporting downstream candidate gene analysis. Within this region, 91 predicted genes from the standard gene set (55,246 genes) were identified and used for downstream analysis (Additional file 3).

**Figure 2.**
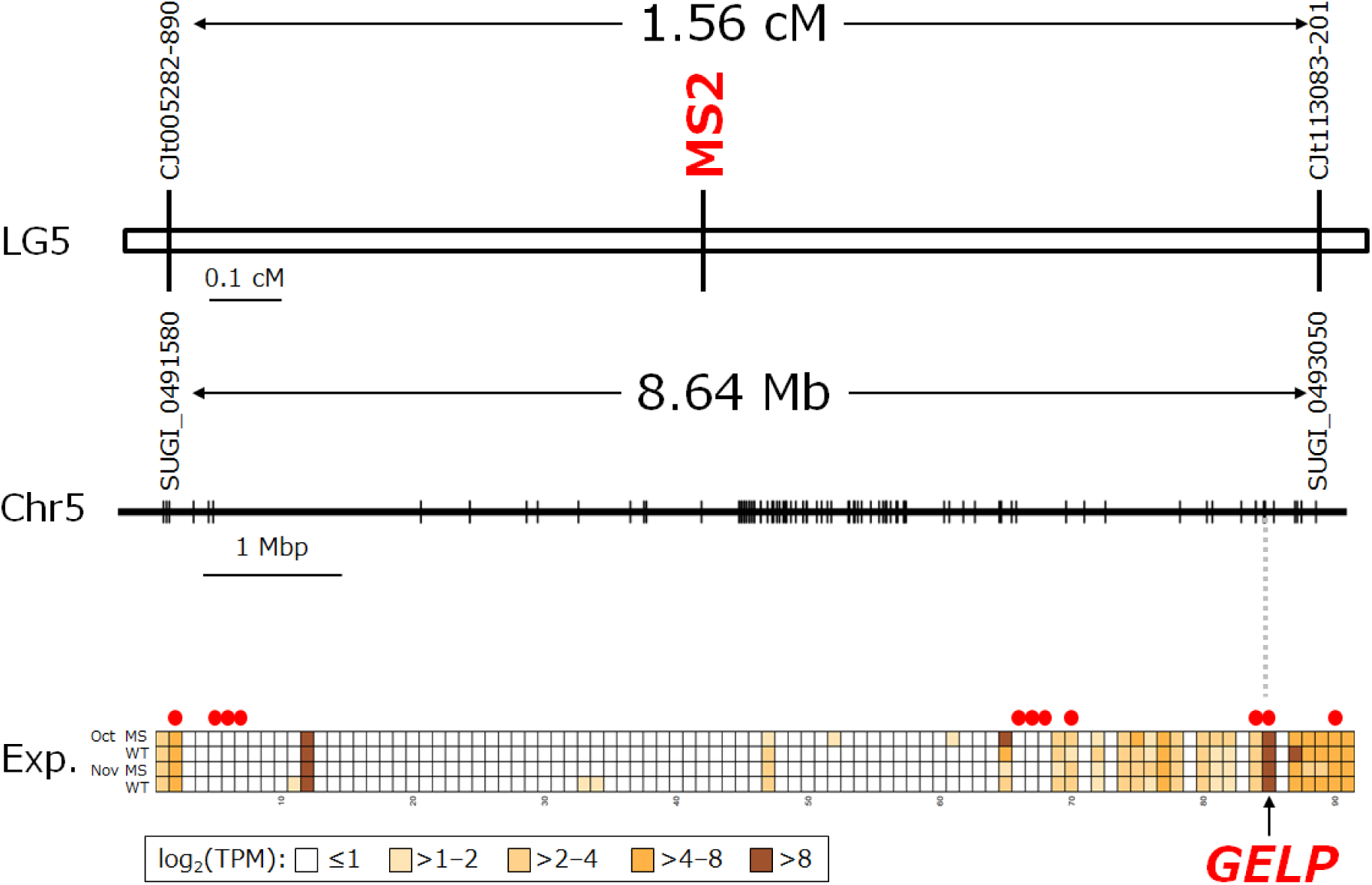
Fine mapping of the MS2 locus and expression profiles of candidate genes. **Top**: Partial linkage map of linkage group 5 (LG5) showing the MS2 locus positioned between two flanking markers. **Middle**: The corresponding 8.64 Mb physical interval on chromosome 5 (Chr5), with vertical lines indicating the positions of 91 positional candidate genes. **Bottom**: Heatmap showing expression levels (TPM) of the 91 genes in male strobili samples collected in October and November from both fertile (WT) and sterile (MS) individuals. Red dots indicate genes functionally annotated as homologs of *Arabidopsis* genes involved in pollen development.

### Identification of candidate genes related to pollen development

To prioritize functionally relevant candidates, we conducted a homology search against 372 *A. thaliana* genes associated with the GO term “pollen development” (GO:0009555), using the TAIR database. A BLASTX search for the nucleotide sequences of *C. japonica* candidate genes and the *Arabidopsis* gene set revealed that 11 out of the 91 MS2-neighboring genes showed significant sequence similarity to pollen development-related genes (Additional file 3).

### Expression profiling in male strobili

To determine whether the candidate genes were expressed in male reproductive organs, RNA-Seq analysis was performed using male strobili collected in October and November from five fertile and three sterile individuals of the S1-2 family (Additional file 1). An average of 7.4 Gb of RNA-Seq data was obtained per sample. The reads were mapped to the SUGI_1 permissive gene set, and transcript abundance was quantified using TPM values. Among the 91 MS2-neighboring genes, 29 genes were expressed (TPM > 1) in at least one of the fertile strobilus samples (Figure 3 and Additional file 3), and all of these genes were in the SUGI_1 standard gene set, ensuring that highly expressed transcripts were not omitted from the expression analysis. Comparison of expression levels between fertile and sterile individuals revealed no statistically significant differences (*p* > 0.01) for these MS2 candidate genes. This suggests that if MS2 is caused by a mutation in one of these genes, it likely does not result from transcriptional suppression in sterile individuals.

**Figure 3.**
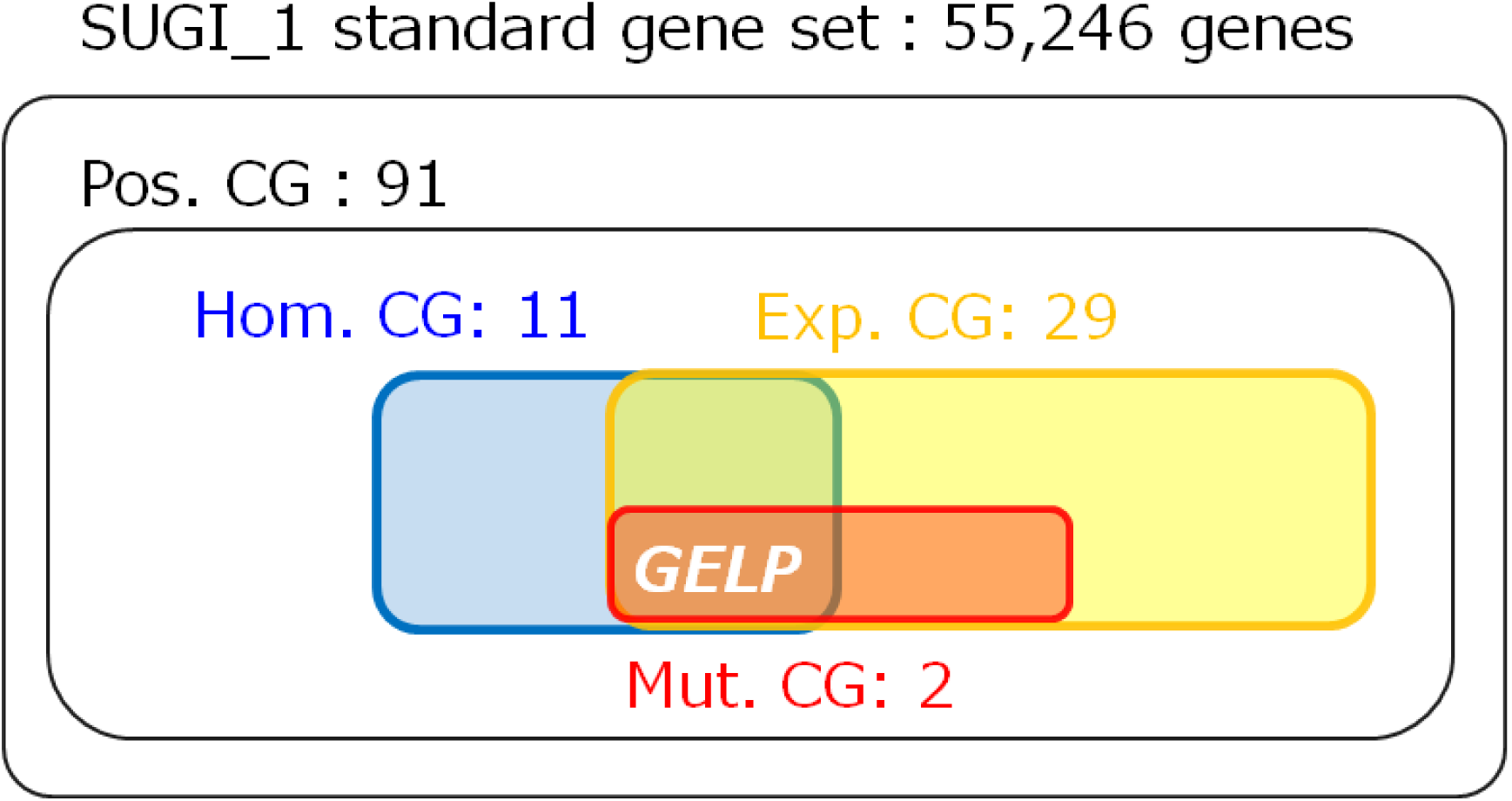
Summary of the candidate gene (CG) screening process, illustrated as a Venn diagram. A total of 91 positional candidate genes (Pos. CG) were initially identified based on genome sequence and linkage analysis. Among these, 11 genes were functionally annotated as homologs of *Arabidopsis* genes involved in pollen development (Homology-based candidate genes; Hom. CG), 29 genes were expressed in male strobili based on RNA-Seq data (Expressional candidate genes; Exp. CG), and 2 genes contained predicted deleterious mutations (Mutational candidate genes; Mut. CG). Only one gene, a GDSL esterase/lipase protein (*GELP*), met all three criteria and was thus identified as the most likely candidate gene responsible for MS2-type male sterility in *C. japonica*. Note: One of the two Mutational CGs (homologous to *Arabidopsis INST1*) does not fall into the Hom. CG category.

### Detection and validation of deleterious mutations

We next examined whether deleterious mutations were present in the coding regions of the 29 expressed genes. Variant calling was conducted using RNA-Seq data, and amino acid substitutions were evaluated using PROVEAN to assess their functional impact. Three deleterious amino acid substitutions (PROVEAN score ≤ –2.5) were identified in two genes (Additional file 3). Notably, one gene (CJt112073 or SUGI_0493010), encoding a GDSL-type esterase/lipase (GELP), contained a substitution from serine to phenylalanine at position 40 (p.S40F) located within the predicted enzymatic active site (Figure 4). The other two mutations were found in a gene homologous to *Arabidopsis INTS1* (SUGI_0492850), which is not known to be involved in pollen development and did not meet the homology-based candidate criteria (Figure 3, Additional file 3). Furthermore, RNA-Seq analysis showed that the *INTS1* mutation was absent in ‘Gosenshi-1’ (Additional file 4), allowing this gene to be excluded from the list of mutational candidates.

**Figure 4.**
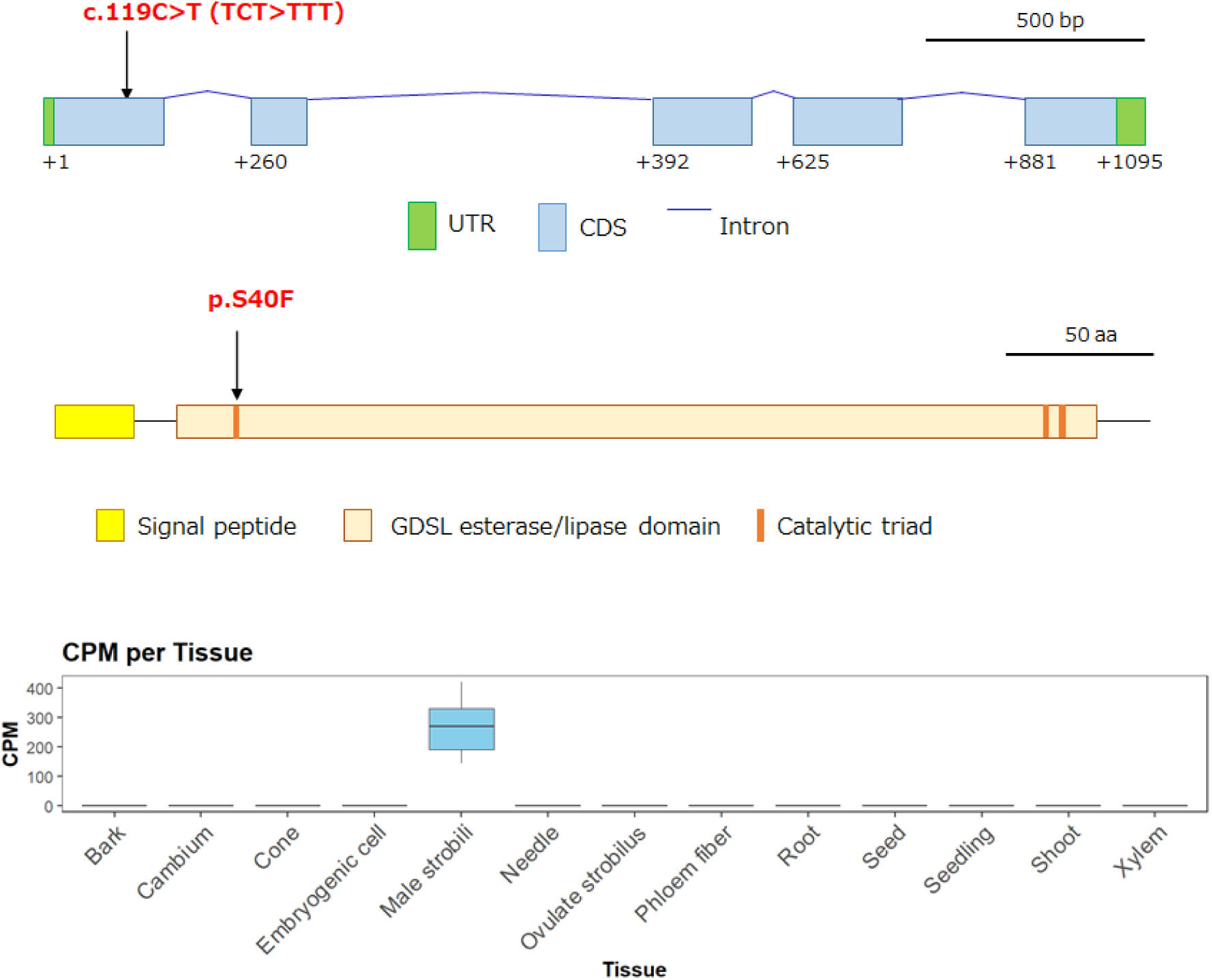
Structural, functional, and expression profiling of the *GELP* gene. **Top**: Exon-intron structure of the *GELP* gene (SUGI_0493010). Coding sequences (CDS) are shown in blue boxes, untranslated regions (UTRs) in green, and introns as lines connecting exons. Numbers bellow the exons indicate the relative positions of the CDS nucleotides. The position of the MS2-associated SNP (c.119C>T, TCT>TTT) is indicated with an arrow, corresponding to a missense mutation in the first exon. **Middle**: Schematic representation of the predicted protein encoded by the *GELP* gene based on InterPro annotation. The signal peptide (yellow), GDSL esterase/lipase domain (light orange), and catalytic triad residues (orange bars) are indicated. The amino acid substitution (p.S40F) caused by the SNP is located within the N-terminal region of the conserved GDSL domain. Only one coding SNP (c.119C>T) leading to the S40F substitution was identified between fertile and sterile haplotypes reconstructed from RNA-Seq data in the S1-2 family. **Bottom**: Expression profile of *GELP* across 13 tissues of C. japonica based on RNA-Seq data from SugiExDB [41]. Counts per million (CPM) values are shown as boxplots. *GELP* exhibits strong and specific expression in male strobili, with no detectable expression in other tissues, including needles, cambium, roots, and ovulate strobili.

To validate the presence of the *GELP* mutation in another genetic background, we examined ‘Gosenshi-1,’ an individual previously confirmed to carry the *MS2* mutation in a heterozygous state (*Ms2*/*ms2*) through controlled crossing experiments [23]. RNA-Seq-based analysis confirmed that the same S40F substitution was present in ‘Gosenshi-1’ (Additional file 4). The consistent presence of this mutation in both *ms2*/*ms2* and *Ms2*/*ms2* individuals supports the hypothesis that *GELP* is the most likely causal gene for MS2-type male sterility in *C. japonica*.

### Structural, functional and expression profiling of GELP

Structural annotation of the *GELP* gene (SUGI_0493010) revealed five exons and four introns (Figure 4, top). TSSPlant predicted multiple transcription start sites (TSSs) within the upstream region of the gene which exceeded the prediction threshold. Among these, one TSS candidate was fully supported by the full-length cDNA sequence obtained by biotinylated CAP trapper method [37]; this site was therefore considered the most likely transcription initiation site. The 5′ untranslated region (UTR) was defined as the 19-bp region between this TSS and the start codon. In contrast, Iso-Seq reads aligned one base upstream of the predicted TSS, likely due to non-templated nucleotide addition by the reverse transcriptase during cDNA synthesis, which can introduce a non-templated nucleotide at the 5′ end [42]. Taken together, these observations support the accuracy of the TSS identified by TSSPlant and the full-length cDNA.

The 3′ end of the transcript was inferred based on the termination pattern of Iso-Seq reads downstream of the stop codon. The majority of reads converged at a position where read depth reached a local minimum, approximately 28 bp downstream of a canonical polyadenylation (polyA) signal (AATAAA), and was followed by a plateau in coverage. This position likely represents the major polyadenylation cleavage site. Additionally, a minor cleavage site was suggested further downstream, approximately 26 bp beyond a secondary polyA signal.

Functional annotation of the predicted protein sequence (364 amino acids) identified a conserved GDSL esterase/lipase domain (PF00657), belonging to the SGNH hydrolase superfamily (IPR036514). The catalytic triad—Ser40, Asp329, and His332—was conserved, and the S40F missense mutation identified in *ms2/ms2* individuals lies within this catalytic region (Figure 4, middle). Notably, the affected serine (S40) is not only a part of the predicted catalytic triad (S40, D329, H332) but also corresponds to the conserved serine residue within the GDSL motif (Gly–Asp–Ser–Leu), which defines this esterase/lipase family. Substitution of this critical serine to phenylalanine (S40F) is therefore expected to abolish enzymatic activity. To confirm the specificity of this mutation, haplotype sequences spanning the GELP coding region were reconstructed from RNA-Seq data for both fertile and sterile individuals in the S1-2 family. Among all identified variants, only a single nucleotide substitution (c.119C>T) corresponding to the S40F amino acid change was found, indicating that this mutation is the sole coding difference between the two haplotypes. These findings strongly support the role of SUGI_0493010 as the causal gene underlying MS2-type male sterility.

In addition to the structural and functional annotations, tissue-specific expression profiles were examined using RNA-Seq data from 13 *C. japonica* tissues available in SugiExDB [41]. CPM values were visually compared across tissues to explore tissue-preferential expression tendencies. GELP exhibited strong and specific expression in male strobili, with no detectable expression in other tissues, including needles, cambium, roots, and ovulate strobili (Figure 4, bottom). This expression pattern supports the involvement of GELP in male reproductive development.

## Discussion

In this study, we identified a strong candidate gene for the MS2-type male sterility in *Cryptomeria japonica* through a comprehensive approach that combined genetic linkage mapping, gene expression analysis, and functional prediction of amino acid substitutions. Our results suggest that a GDSL-type esterase/lipase gene (*GELP*), which harbors a deleterious amino acid substitution, is the most likely causal gene for MS2-type male sterility.

The candidate *GELP* gene resides within the 1.56 cM interval surrounding the *MS2* locus on linkage group 5, corresponding to a 8.64 Mb region of the reference genome. Among the 91 genes in this interval, *GELP* was one of 11 genes with homology to *Arabidopsis* genes involved in pollen development and one of 29 genes expressed in male strobili. Expression analysis showed that *GELP* is specifically expressed in male strobili (Figure 4), and is present in both fertile and sterile individuals (Figure 2). This supports the idea that the sterility phenotype arises not from a lack of expression, but from a functional disruption caused by the amino acid substitution. Notably, the gene exhibited a serine-to-phenylalanine substitution (S40F) within a predicted active site (Figure 4), which was predicted by PROVEAN to be functionally deleterious. Although the deleterious effect of the S40F substitution was computationally predicted by PROVEAN, the affected serine residue is part of the conserved catalytic triad of GDSL esterases/lipases. This serine hydroxyl group is essential for nucleophilic attack during catalysis, and its substitution with a bulky, nonpolar phenylalanine is expected to abolish enzymatic activity. Furthermore, the same mutation was identified (Additional file 4) in a heterozygous (*Ms2*/*ms2*) individual, ‘Gosenshi-1,’ which was derived from an independent genetic background, indicating that this mutation is consistently associated with MS2-type male sterility in *C. japonica*.

GDSL esterases/lipases have been implicated in a variety of biological processes, including lipid metabolism, cuticle formation, and reproductive development in plants [43–46]. In *Arabidopsis thaliana*, mutations in orthologous genes have been shown to cause male sterility due to defects in pollen wall formation [47]. Given the role of sporopollenin and lipids in pollen wall structure, it is plausible that the *GELP* gene in sugi plays a similar role in pollen development. Supporting this hypothesis, several morphological characteristics observed in *ms2* mutant individuals are consistent with those reported in *A. thaliana*. Loss-of-function mutations in orthologous GDSL genes result in abnormal pollen wall formation, irregularly shaped microspores, early microspore degeneration, and partial or complete nuclear loss [46]. Notably, clumping or high-density aggregation of microspores— likely due to defects in post-meiotic microspore separation—has also been observed in MS2-type male sterility in *C. japonica*, where densely packed and irregular microspores, some lacking nuclei, are found [9, 10]. These shared phenotypes across species strongly support the idea that GELP plays a conserved and essential role in male gametophyte development, likely through contributing to proper microspore maturation.

Despite this collective evidence supporting *GELP* as the candidate gene, several questions remain to be resolved. Microscopic observations revealed that some individuals classified as sterile based on genetic and phenotypic data still produced visible pollen grains (Additional file 5), confirming that the *ms2*/*ms2* genotype may not always result in complete pollen abortion [10]. One possible explanation is the presence of a neighboring gene, SUGI_0493000, which encodes another GDSL esterase/lipase protein and is located adjacent to *GELP* (SUGI_0493010). Although the expression level of SUGI_0493000 is relatively low (Figure 2 and Additional file 3), it may exert a weak compensatory effect under certain conditions, thereby allowing partial pollen development in *ms2/ms2* individuals. Alternatively, this inconsistency may be due to environmental influences, measurement variability, or the presence of modifier loci (small-effect QTLs) [48–50]. Furthermore, the segregation ratio of fertile and sterile individuals in the S1-2 family deviated from the expected 1:1 Mendelian ratio. This may indicate viability effects or segregation distortion related to the MS2 locus, possibly due to impaired pollen function or reduced viability of early *ms2*/*ms2* embryos. These observations highlight the need for further studies to clarify the penetrance and expressivity of the *MS2* mutation.

To definitively confirm the causal role of the *GELP* gene in MS2-type male sterility, functional validation is necessary. Genome editing technologies such as CRISPR/Cas9, which have recently been successfully applied to *C. japonica* [51, 52], could be employed to introduce or correct the identified mutation. This would allow direct assessment of the gene’s role in pollen development and provide a powerful tool for breeding programs aimed at producing pollen-free sugi trees. Additionally, broader surveys across natural and breeding populations may reveal further allelic diversity at the MS2 locus, similar to findings from our previous study [17] that analyzed the MS1 locus using amplicon sequencing and haplotype network construction.

The identification of *GELP* as a candidate gene for MS2-type male sterility represents a significant step toward understanding the molecular basis of male sterility in *C. japonica* and developing effective strategies for reducing pollen production. This research contributes to both fundamental knowledge of conifer reproduction and practical efforts to mitigate the growing public health issue of cedar pollen allergy.

## Conclusions

In this study, we identified a strong candidate gene for *MALE STERILITY 2* (*MS2*) in *Cryptomeria japonica* using a map-based approach that integrated high-resolution linkage mapping, transcriptome profiling, and mutation analysis. The *MS2* locus was localized to a 8.64 Mb region on chromosome 5, within which a GDSL-type esterase/lipase gene (*GELP*) was identified as the most likely causal gene. A deleterious amino acid substitution (S40F) was detected in sterile individuals and validated in an independent heterozygous line, supporting the association between the mutation and the *MS2* sterility phenotype. These findings provide a key genetic resource for breeding pollen-free *C. japonica* trees, which are of increasing social importance in the context of allergenic pollen mitigation. Moreover, the identification of *GELP* as a candidate gene adds to our understanding of the molecular mechanisms underlying male sterility in conifers and highlights the utility of integrative genomic approaches in forest tree research.

## List of abbreviations

CDS: Coding Sequence
DEG: Differentially Expressed Gene
GELP: GDSL esterase/lipase protein
GO: Gene Ontology
INTS1: Integrator Complex Subunit 1
kb: kilobase pairs
LG: Linkage Group
Mb: megabase pairs
MS: Male Sterility
QTL: Quantitative Trait Locus
RNA-Seq: RNA Sequencing
SNP: Single Nucleotide Polymorphism
TPM: Transcripts Per Million
TSS: Transcription Start Site
UTR: Untranslated Region.

## Declarations

### Ethics approval and consent to participate

Not applicable.

### Consent for publication

Not applicable.

### Availability of data and materials

The RNA-Seq Illumina reads were deposited in the Sequence Read Archive database of the DNA Data Bank of Japan (DDBJ) under accession number DRR733425–DRR733440. All additional datasets and materials used during the current study are available from the corresponding author upon reasonable request.

### Competing interests

The authors declare that they have no competing interests.

### Funding

This work has been supported in part by JSPS KAKENHI Grant Numbers JP16H06279 (PAGS), JP20H03239, JP23K26956 and JP23H02263, the Ministry of Agriculture, Forestry and Fisheries’ “Agriculture, Forestry and Fisheries and Food Industry Science and Technology Research Promotion Project” and the research program on development of innovative technology grants (JPJ007097) from the Project of the Bio-oriented Technology Research Advancement Institution (BRAIN) (Project ID 28013BC), and NIBB Collaborative Research Program (15-829, 16-403, 17-405, 18-408, 19-420, 20-428, 21-302, 22NIBB402, and 23NIBB405).

### Authors’ contributions

SU and YM conceived and designed the study. YH and MT performed the experiments. YH conducted the linkage analysis. HK proposed the strategy for the phenotypic evaluation of male strobili. SU performed the bioinformatic analyses. KY, SS, and YS generated early sequencing datasets. TF and MK assembled the draft genome, which served as the reference for the initial linkage mapping. TU contributed to discussions and provided critical comments throughout the research process. JI and SH established the mapping population and provided the breeding material. SU wrote the initial draft of the manuscript with input from all authors. All authors read and approved the final manuscript.

## Acknowledgements

A part of bioinformatic analysis was carried out by the supercomputer of AFFRIT, MAFF, Japan. The Arboretum and Nursery Office, FFPRI was involved in the maintenance of the *C. japonica* nursery. We thank Eriko Tsurisaki and Nana Matsumura of Niigata University for photographs of male strobili and light microscopic images for wild-type and male-sterile *C. japonica*. Yasuyuki Komatsu and Nozomi Ohmiya, FFPRI, performed DNA and RNA extraction, respectively. ChatGPT (OpenAI) was used to suggest improvements in grammar and phrasing during the writing process. All scientific content, data analysis, and interpretations were solely developed and validated by the authors.

## Additional files

### Additional file 1

File name: Additional file 1

File format: .xlsx

Title of data: **Supplementary Table 1** RNA-Seq sample information

Description of data: List of sample ID, collection date, library ID, sequencing depth, and fertility status.

### Additional file 2

File name: Additional file 2

File format: .xlsx

Title of data: **Supplementary Table 2** Details of markers and the fine mapping

Description of data: Marker IDs and their primer sequences; Draft genome contig IDs of marker origin; SUGI_1 (chromosome, SNP position and gene IDs) of marker origin; linkage map positions (cM); and genotyping results for 129 individuals in S1-2 family. Marker IDs beginning with ‘AX-’ are used in our previous linkage map [22], and their genotypes were compiled from <https://doi.org/10.1371/journal.pone.0206695.s009>.

### Additional file 3

File name: Additional file 3

File format: .xlsx

Title of data: **Supplementary Table 3.** List of genes located on MS2 locus, and their characteristics Description of data: Genes located around MS2 locus around 1.56 cM, functional similarity to *Arabidopsis thaliana* genes, TPM values, and non-synonymous amino acid mutations with its PROVEAN score. In addition to the homology search against *Arabidopsis* pollen-related genes (TAIR GO:0009555) described in the Methods section, previously obtained functional annotations based on the full ARAPORT11 protein from our earlier study [26] are included for reference. These supplementary annotations were not used for candidate gene selection but are provided here to assist interpretation. The functional annotation dataset is available at https://forestgen.ffpri.go.jp/en/info_sugi1.html.

### Additional file 4

File name: Additional file 4

File format: .pdf

Title of data: **Supplementary Figure 1**. Amino-acid sequence alignment of the *MS2* mutational candidate genes in *Cryptomeria japonica*: (A) *INTS1* and (B) *GELP*.

Description of data: Alignment of the amino acid sequences of SUGI *standard gene set* (SUGI_0492850 and SUGI_0493010 for *INTS1* and *GELP*, respectively), Gosenshi-1_a1 (wild-type allele), Gosenshi-1_a2 (mutant allele), and the *ms2* mutant from the S1-2 family, generated in CLC Genomics Workbench ver. 20.0.4 (Qiagen). Residues are colored according to side-chain polarity. Red arrows indicate positions of deleterious amino acid substitutions predicted by PROVEAN (score ≤ –2.5). For INTS1, two deleterious amino acid substitutions (at positions 613 and 1047) were identified in the S1-2 family, but these substitutions were absent in ‘Gosenshi-1.’ For GELP, two substitutions are observed relative to SUGI_0493010 (S40F and C352Y), but only S40F is deleterious and uniquely distinguishes the *ms2* mutant from the wild-type allele. In addition, the 352nd residue in ‘Gosenshi-1’ (*Ms2*/*ms2*) is homozygous for tyrosine (Y/Y), as indicated by the blue arrow, further indicating that C352Y is unlikely to be the causal mutation.

### Additional file 5

File name: Additional file 5

File format: .pdf

Title of data: **Supplementary Figure 2** Microscopic cross-section of a male strobilus from a sterile individual (Y677) in S1-2 family

Description of data: A partial formation of pollen was observed in the male strobilus of an individual with the *ms2/ms2* genotype. The sample was collected from the Chiyoda nursery on 20 December 2023. The scale bar is based on an approximate size of the male strobilus (∼4 mm) and is provided as a reference only.

## Notes

### Competing Interest Statement

The authors have declared no competing interest.

