## Supplementary figures and images for "Positional identification of a candidate gene for *MALE STERILITY 2* (*MS2*) by linkage mapping and transcriptomic data in *Cryptomeria japonica* D. Don"

### Additional file 4

(A) *INTS1*

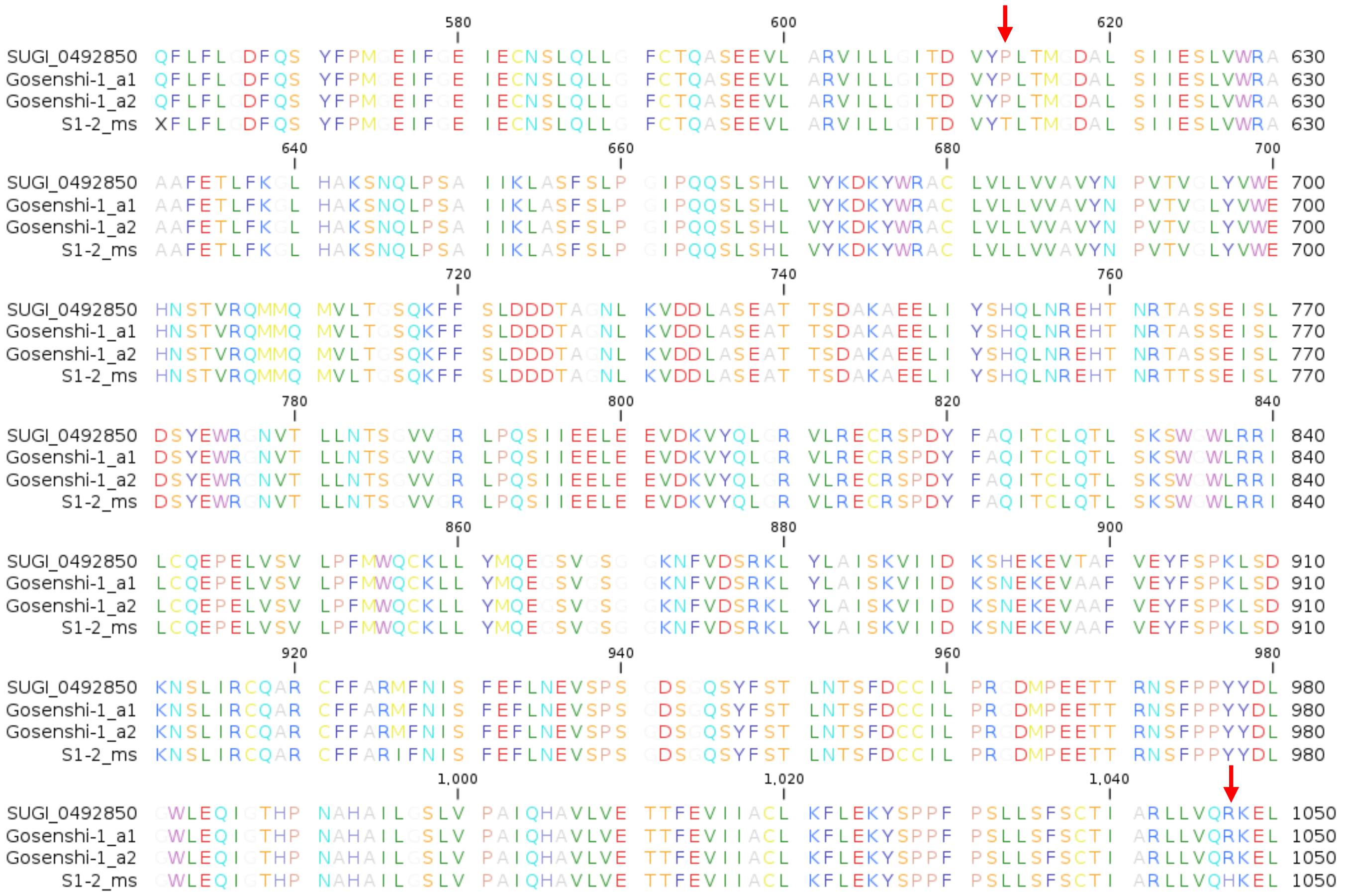


(B) *GELP*

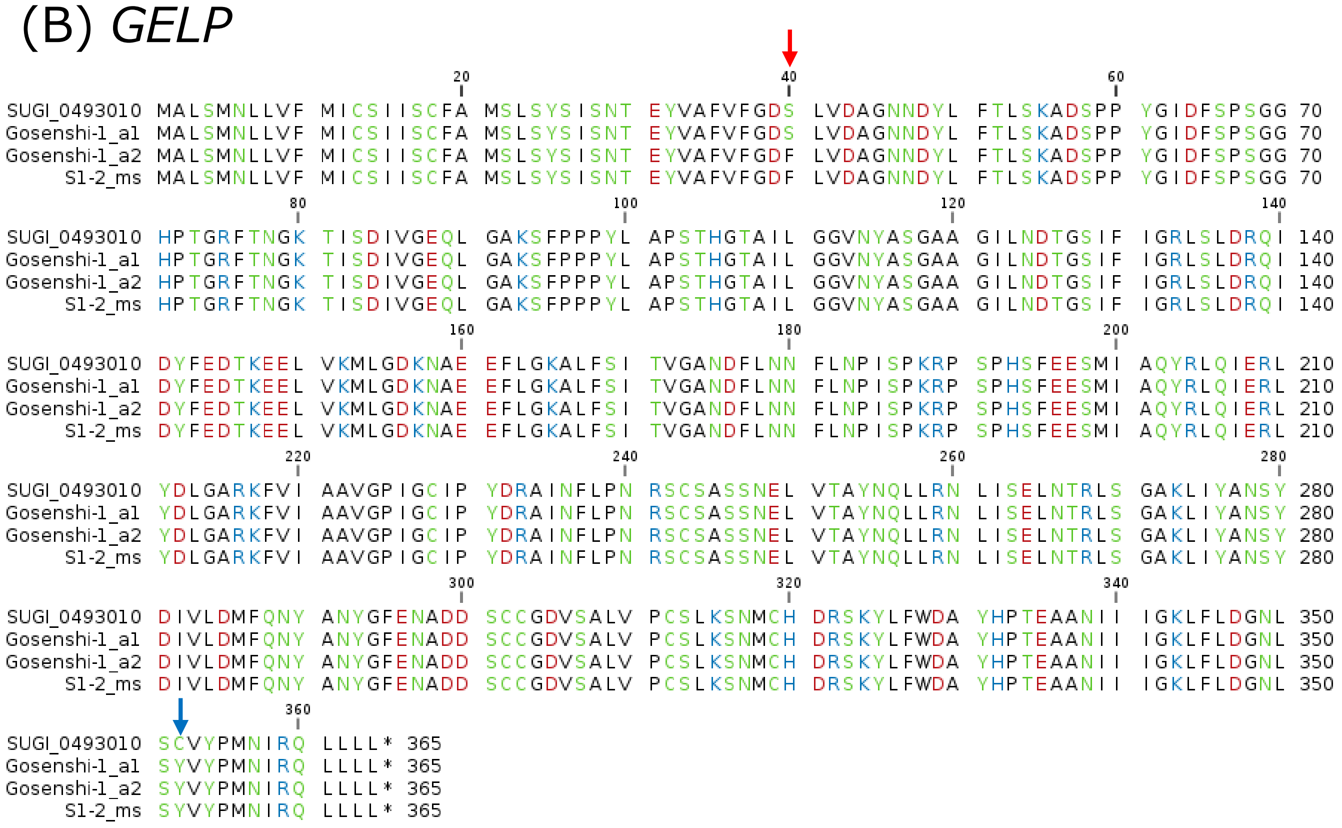

### Additional file 5

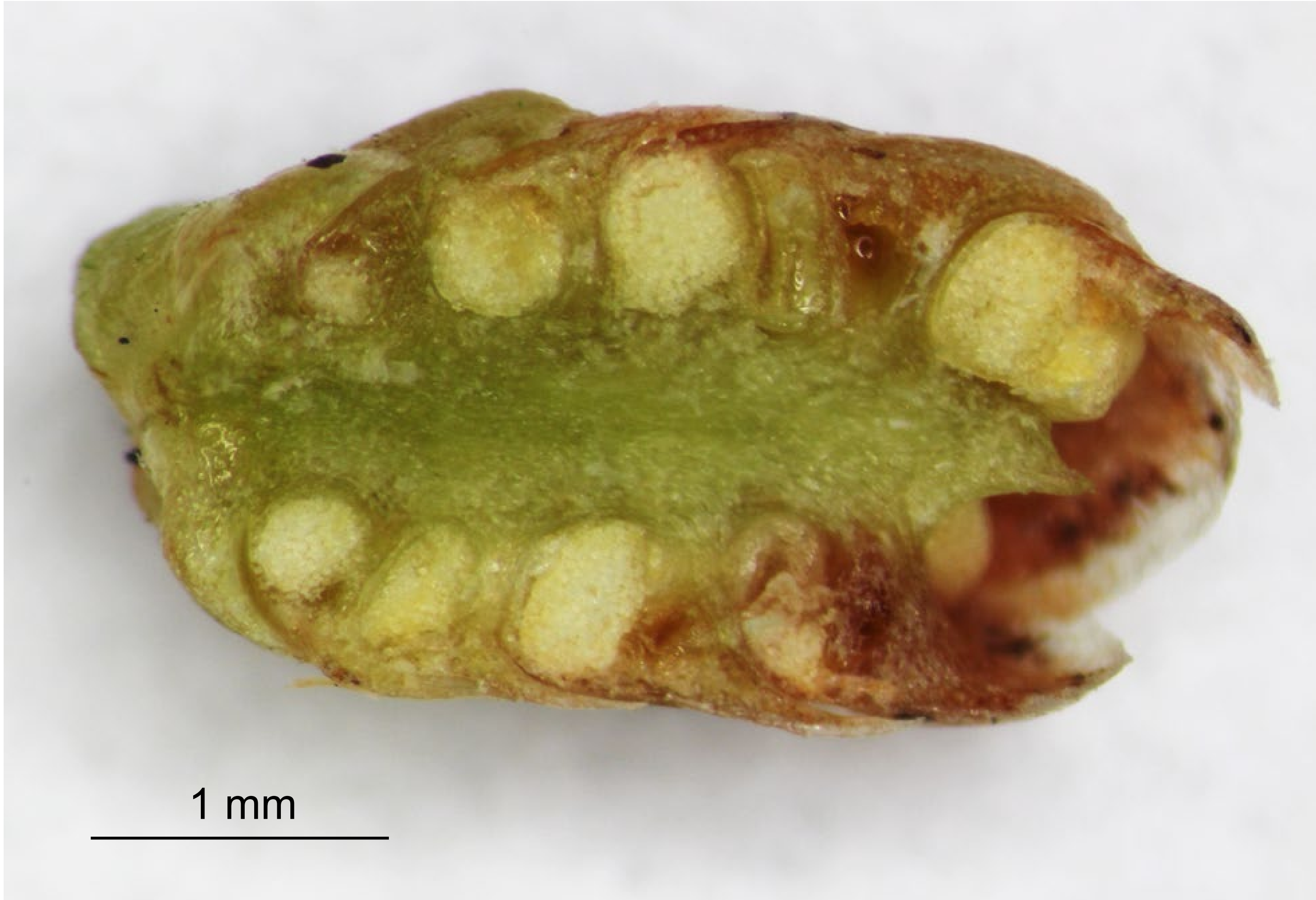
